# Defining marine microbial biomes from environmental and dispersal filtered metapopulations

**DOI:** 10.1101/141077

**Authors:** Markus V. Lindh

## Abstract

Energy and matter fluxes essential for all life^1^ are modulated by spatial and temporal shifts in microbial community structure resulting from environmental and dispersal filtering^2,3^, emphasizing the continued need to characterize microbial biogeography^4,5^. Yet, application of metapopulation theory, traditionally used in general ecology for understanding shifts in biogeographical patterns among macroorganisms, has not been tested extensively for defining marine microbial populations filtered by environmental conditions and dispersal limitation at global ocean scales. Here we show, from applying metapopulation theory on two major global ocean datasets^6,7^, that microbial populations exhibit core- and satellite distributions with cosmopolitan compared to geographically restricted distributions of populations. We found significant bimodal occupancy-frequency patterns (the different number of species occupying different number of patches) at varying spatial scales, where shifts from bimodal to unimodal patterns indicated environmental and dispersal filtering. Such bimodal occupancy-frequency patterns were validated in Longhurst’s classical biogeographical framework and *in silico* where observed bimodal patterns often aligned with specific biomes and provinces described by Longhurst and where found to be non-random in randomized datasets and mock communities. Taken together, our results show that application of metapopulation theory provides a framework for determining distinct microbial biomes maintained by environmental and dispersal filtering.

---

Discovery of biogeographical patterns in marine microbial assemblages^6,7,9–12^ caused considerable excitements because it indicated distribution patterns analogous to macroecological patterns^13–17^. These observations have provided clues that marine microbes are, at least to some extent, limited by the environment and their dispersal capacity rather than being cosmopolitan.

Metapopulation theory is a key framework in general ecology and incorporates population dispersal to empirically test occupancy-frequency distributions of different organisms ranging from insects to plants^18^. Predictions of occupancy-frequency patterns have been developed in e.g. the core- and satellite hypothesis (CSH) by Ilkka Hanski^19^ where the so-called rescue effect supports the survival of metapopulations, and forms a bimodal occupancy-frequency pattern (See Fig. 1A). Previous results applying the CSH to marine ecosystems suggest that this approach could potentially allow for precise definitions of microbial biomes^20^.

**Figure 1.**
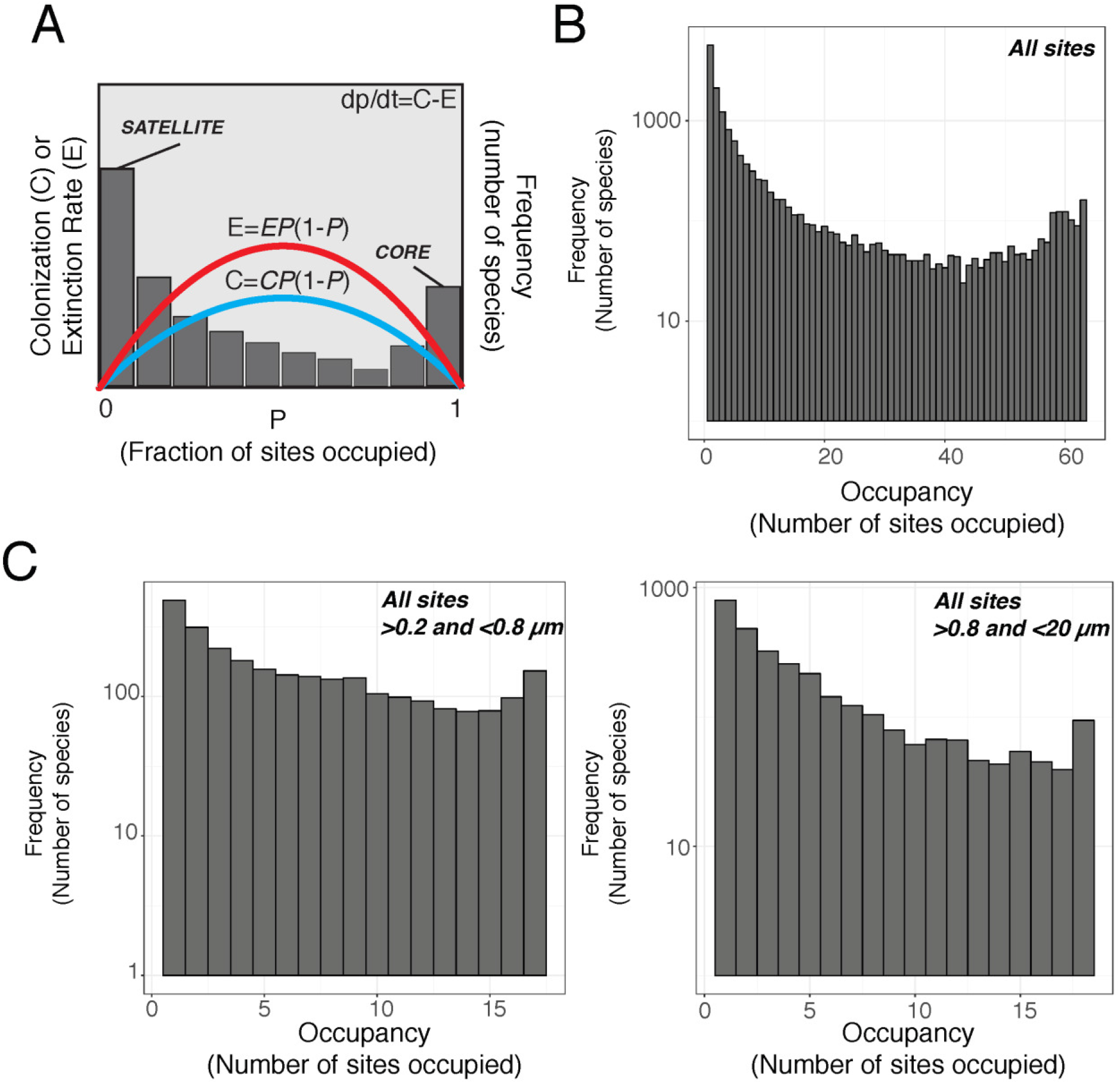
Conceptual drawing of the core satellite hypothesis (CSH; A), and Occupancy-frequency distributions (the different number of populations occupying different number of sites) of populations estimated from 16S rRNA gene amplicon and metagenomic data obtained from the Tara oceans^6^ (B), and Malaspina^7^ (C) datasets. (A) was modified from^20^; *P* is the fraction of occupied sites, *C* is colonization rate and *E* is extinction rate. The quadratic colonization and extinction rates are calculated according to d*P*/d*t* = *CP*(1 – *P*) – *EP*(1 – *P*)^19^. The CSH predicts a bimodal pattern, and incorporates positive feedback mechanisms between local abundance and regional occupancy^19^. High rates of colonization in the CSH protect a population from extinction and are known as the rescue effect.

Here we collected data on bacterial and archaeal populations (estimated from 16S rRNA gene sequencing and metagenomic data) obtained from two recent major global ocean datasets, Tara oceans^6^ and Malaspina^7^ (Fig. S1) to examine the shape of occupancy-frequency distributions in global ocean data and aimed to define distinct microbial biomes.

For the Tara oceans dataset^6^ we focused our analyses on surface seawater (≤5 m) and analyzed 63 stations in total (Fig. S1). Occupancy-frequency analysis for the whole transect revealed a significant bimodal pattern, with the number of populations demonstrating a monotonic decrease with increasing number of sites occupied followed by an increase in the number of populations occupying all sites (Fig. 1B; Table 1; Mitchell-Old’s and Shaw’s test, p<0.05). In addition, populations affiliated with Alphaproteobacteria (represented mainly by the SAR11 clade bacteria) also exhibited bimodality for the whole transect (Table 1; Mitchell-Old’s and Shaw’s test, p<0.05). In contrast, Euryarchaeota, Cyanobacteria, Gammaproteobacteria and Bacteroidetes did not exhibit significant bimodal patterns (Table 1; Mitchell-Old’s and Shaw’s test, p>0.05). Such results indicate that, among individual taxa, except Alphaproteobacteria, surface ocean microbial communities can be limited by the prevailing environmental conditions and their dispersal capacity.

**Table 1.** Prevalence of bimodal compared to unimodal occupancy-frequency patterns within the Tara Oceans and Malaspina transects for different oceanic basins and within Longhurst’s biomes and provinces. Asterisks denote level of significance (*** <0.001, ** 0<.01, *<0.05).

| Dataset | Region | Taxa |  |  |  |  |  |
| --- | --- | --- | --- | --- | --- | --- | --- |
|  |  | ALL OTUs<br>(n=35651) | Euryarchaeota<br>(n=550) | Cyanobacteria<br>(n=963) | Alphaproteobacte<br>ria (n=7151) | Gammaproteobacteri<br>a (n=7766) | Bacteroidetes (n=3380) |
| Tara Oceans | Full transect | BIMODAL** | OTHER | OTHER | BIMODAL*** | OTHER | OTHER |
|  | South Pacific | BIMODAL*** | OTHER | BIMODAL* | BIMODAL*** | BIMODAL*** | BIMODAL*** |
|  | North Atlantic | BIMODAL*** | BIMODAL*** | BIMODAL*** | BIMODAL*** | BIMODAL** | BIMODAL** |
|  | Coastal Biome (L) | BIMODAL* | OTHER | OTHER | BIMODAL*** | BIMODAL* | OTHER |
|  | Trades Biome (L) | BIMODAL*** | OTHER | BIMODAL* | BIMODAL*** | BIMODAL*** | BIMODAL*** |
|  | Westerlies (L) | BIMODAL*** | OTHER | BIMODAL* | BIMODAL*** | BIMODAL*** | BIMODAL*** |
|  | MEDI (L) | BIMODAL*** | UNIMODAL | BIMODAL* | BIMODAL*** | BIMODAL*** | BIMODAL*** |
|  | EAFR (L) | BIMODAL*** | BIMODAL*** | BIMODAL*** | BIMODAL*** | BIMODAL** | BIMODAL*** |
|  | SATL (L) | BIMODAL** | UNIMODAL | BIMODAL*** | BIMODAL*** | UNIMODAL | BIMODAL* |
|  | SPSG (L) | BIMODAL*** | OTHER | BIMODAL* | BIMODAL*** | BIMODAL*** | BIMODAL*** |
|  | NASW (L) | BIMODAL*** | BIMODAL*** | BIMODAL*** | BIMODAL*** | BIMODAL* | BIMODAL* |

**Table 1** continued.
| Dataset | Region | Taxa |  |  |  |  |  |
| --- | --- | --- | --- | --- | --- | --- | --- |
|  |  | ALL OTUs (n=3902) | Thaumarchaeota<br>(n=48) | Gammaproteobacteria<br>(n=384) | Alphaproteobacteria<br>(n=405) | Actinobacteria (n=143) | Deltaproteobacteria<br>(n=433) |
| Malaspina<br>0.2/0.8 µm | Full transect | BIMODAL*/ BIMODAL | OTHER/OTHER | OTHER/OTHER | OTHER/BIMODAL*** | OTHER/OTHER | OTHER/OTHER |
|  | South Atlantic | BIMODAL**/BIMODAL* | OTHER/BIMODAL** | BIMODAL**/BIMODAL** | OTHER/BIMODAL** | UNIMODAL/UNIMODAL | BIMODAL*/UNIMODAL |
|  | Indian Ocean | ND/BIMODAL* | ND/OTHER | ND/BIMODAL** | ND/BIMODAL** | ND/BIMODAL*** | ND/UNIMODAL |
|  | WMbSC 2: NADW–CDW–AABW | BIMODAL*/UNIMODAL | BIMODAL***/ND | BIMODAL**/ND | BIMODAL**/ND | OTHER/UNIMODAL | BIMODAL*/UNIMODAL |
|  | WMbSC 4: Purest AABW Ross | ND/BIMODAL* | ND/BIMODAL*** | ND/BIMODAL*** | ND/BIMODAL*** | ND/BIMODAL* | UNIMODAL/BIMODAL* |
|  | SATL (L) | UNIMODAL/BIMODAL* | OTHER/OTHER | BIMODAL***/BIMODAL* | OTHER/BIMODAL*** | OTHER/BIMODAL* | UNIMODAL/UNIMODAL |

For subsets in the whole Tara Oceans transect, here exemplified by the South Pacific and North Atlantic, we found significant bimodal occupancy-frequency patterns for the total community (Table 1; Mitchell-Old’s and Shaw’s test, p<0.05). We note that Euryarchaeota populations only exhibited a bimodal pattern in the North Atlantic basin (Table 1; Mitchell-Old’s and Shaw’s test, p<0.05). Our results therefore indicate that the total community and most individual taxa are bimodal within a coherent oceanic region. Reciprocally, particular microbial groups are subjected to environmental and dispersal filtering^21^ at different spatial scales resulting in geographically constricted populations.

The Malaspina dataset contained 30 stations collected from the deep-sea, typically ≥4000 m deep (Fig. S1). This dataset also included size-fractionated samples corresponding to free-living (>0.2 and <0.8 μm) and particle-attached (>0.8 and <20 μm) microbial assemblages. Analysis of occupancy-frequency patterns among deep-sea microbial assemblages revealed a significant bimodal pattern of the total community in the whole transect for both free-living and particle-attached populations (Fig. 1C; Table 1). Thus, although most deep-sea microbes have been suggested to be confined to a specific region, our results indicate that a core deep-sea community are still distributed across the whole Malaspina transect, suggesting a similar microbial biome without environmental or dispersal filtering.

Variations in the shape of occupancy-frequency distributions in the Malaspina transect were found for different size fractions, taxa, oceanic basins and taxa within different basins (Table 1). For example, Thaumarchaeota populations displayed bimodality in the South Atlantic in the particle-attached community (Table 1; Mitchell-Old’s and Shaw’s test, p<0.05) but not in the free-living community (Table 1; Mitchell-Old’s and Shaw’s test, p>0.05). In the South Atlantic, Gammaproteobacteria populations exhibited a bimodal pattern in both the particle-attached and free-living community (Table 1; Mitchell-Old’s and Shaw’s test, p<0.05). We note that samples collected within the same water mass, here exemplified by “WMbSC 2: NADW-CDW-AABW” and ”WMbSC 4: Purest AABW Ross”, often showed a bimodal occupancy-frequency pattern (Table 1; Mitchell-Old’s and Shaw’s test, p<0.05). Overall these findings substantiate that particular taxa such as Thaumarchaeota can be limited by environmental conditions and dispersal capacity and thus geographically restricted.

We further aimed to validate our observed microbial biomes estimated from bimodal occupancy-frequency patterns in the framework of Longhurst’s ecological biomes and provinces that define biogeographic distributions derived in large part from satellite-based estimations of sea-surface chlorophyll *a* (Chl *a*) concentrations. Longhurst defines four primary oceanic biomes; the Westerlies, Trades, Polar and Coastal (Fig. S1). These biomes differ in nutrient supply, light and seasonal variation in water column mixing. The biomes are in turn subdivided into several provinces based on Chl *a* distributions (Fig. S1). Our analysis revealed that samples collected as part of the Tara Oceans transect within the Coastal, Westerlies and Trades biomes displayed a distinct bimodal occupancy-frequency pattern (Table 1; Mitchell-Old’s and Shaw’s test, p<0.05). In addition, provinces such as the Indian Monsoon Gyres Province (MONS) and South Atlantic Gyre Province (SATL), exhibited significant bimodal patterns (Table 1; Mitchell-Old’s and Shaw’s test, p<0.05).

During the Malaspina transect examining the deep-sea microbes, only the particle-attached microbial community exhibited bimodal occupancy-frequency in the SATL (Table 1; Mitchell-Old’s and Shaw’s test, p<0.05). Gammaproteobacteria displayed bimodality in the SATL both within the free-living and particle-attached community (Table 1; Mitchell-Old’s and Shaw’s test, p<0.05). However, only particle-attached Alpha- and Actinobacteria displayed bimodality in SATL (Table 1; Mitchell-Old’s and Shaw’s test, p<0.05). Collectively, these findings emphasize that occupancy-frequency patterns among microbial assemblages can likely be used to validate and significantly extend analyses of oceanic divisions as determined by e.g. Chl *a* satellite data^8^.

The difficulty in assessing and discerning sampling effects from true biological effects for understanding metapopulation dynamics is well recognized in terrestrial studies of macroorganisms^18^. Here we aimed to (i) examine the effect of sequencing depth on the cumulative number of core and satellite populations, and (ii) elucidate if bimodal patterns could arise from pure chance and hence be an artefactual effect in marine microbial assemblages. The number of core and satellite populations in subsampled rarefied datasets reached saturation around 25,000 sequence reads (Fig. S2). *In silico* tests with randomized datasets and mock communities confirmed that bimodal patterns were non-random, as no significant bimodal pattern was found in any randomized dataset or mock communities (Fig 2A-B; Mitchell-Old’s and Shaw’s test, p>0.05). Our extensive *in silico* analyses provided a first clue that bimodality could be a biological pattern rather than an artefactual effect.

**Figure 2.**
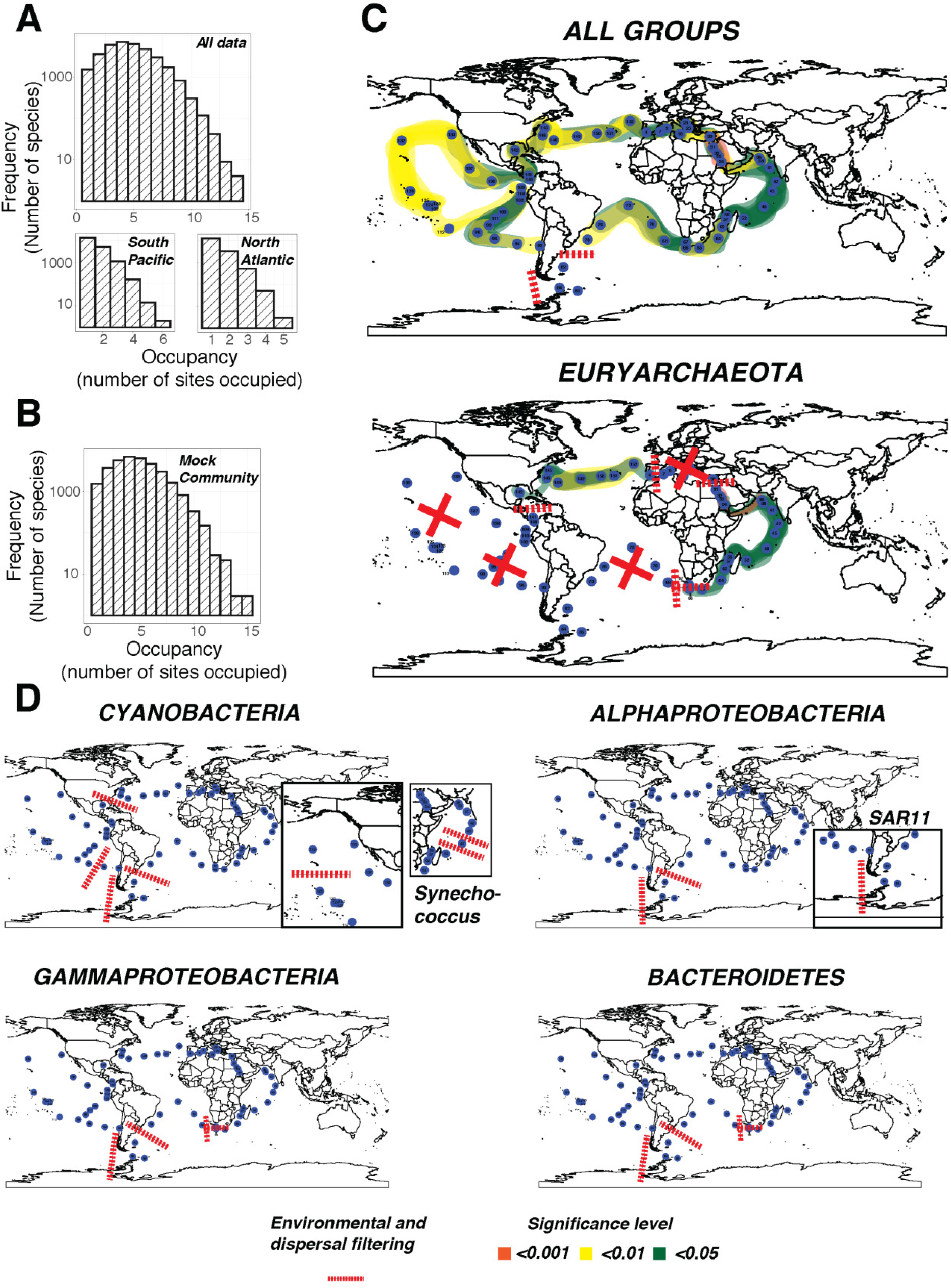
*In silico* tests of randomized sequence data and microbial biomes defined by shifts from bimodal occupancy-frequency patterns to unimodal patterns. Randomizations of observational data were performed for complete datasets and specific oceanic regions (A), and in mock communities (B). The typical occupancy-frequency pattern of the randomized datasets was characterized by most populations being detected at 2-4 sites with a monotonical decrease in the number of populations occupying increasing number of sites. No randomized community dataset exhibited a significant bimodal pattern. The randomized mock community typically displayed prevalence of most populations occupying 2-4 sites but with no significant bimodal pattern found. Microbial biomes (C-D) were defined by shifts from bimodal occupancy-frequency patterns to unimodal patterns by testing the occupancy-frequency distributions with combinations of sites within the Tara oceans transect. Red dashed lines denote environmental and dispersal filtering of microbial assemblages resulting geographically constricted populations. Color denotes significance level of Mitchell-Old’s and Shaw’s test for bimodality.

To test the possibility of allowing the metapopulation framework to pinpoint microbial biomes without testing against a pre-defined oceanic region we analyzed 5 patches at a time in sequence along the Tara oceans transect (Fig. 2C-D). This exercise highlighted oceanic regions where the core populations (bimodal occupancy-frequency patterns; Mitchell-Old’s and Shaw’s test, p<0.05) prevailed and the community was not limited by the environment or dispersal capacity. Reciprocally, we could also define specific microbial biomes along the transect (non-bimodal occupancy-frequency patterns; Mitchell-Old’s and Shaw’s test, p>0.05). It was noteworthy that for all taxa combined, stations obtained in the Southern Ocean limited the distribution of microbial assemblages (Fig. 2C).

For individual taxa in this unsupervised model, the occupancy-frequency pattern varied substantially and different major microbial groups were subjected to different environmental and dispersal filters. Euryarchaeota only had significant bimodal patterns in patches sampled from the North Atlantic, Red Sea, Arabian Sea and the Indian Ocean. Cyanobacteria exhibited bimodality in the Pacific Ocean but the pattern broke down upon entry into the Atlantic Ocean (Fig. 2D). Notably, within Cyanobacteria, *Synechococcus* had a geographical restriction between the North and South Pacific Ocean (see insert Fig. 2D). *Synechococcus* were also limited in their distributions in the Indian Ocean by a single site. Alphaproteobacteria had a similar metapopulation distribution limitation as the total community and Cyanobacteria in the Southern Ocean. Yet, within Alphaproteobacteria, SAR11 clade bacteria (see insert Fig. 2D) had a wider distribution, exemplified by a bimodal pattern between sites obtained in the South Atlantic and Southern Ocean. Still, SAR11 were limited upon entry into the Pacific Ocean. Gammaproteobacteria and Bacteroidetes displayed a clear distinction between samples obtained in the Indian Ocean and South Atlantic where the core community was constrained by the station outside Cape Town.

Taken together, we demonstrate that empirical tests of metapopulation dynamics allows for biogeographical analyses of marine microbes to define microbial biomes. We note that a sequence depth of 25,000 sequence reads is sufficient to capture most of the core- and satellite populations and could be considered as lower limit for microbial biogeography analyses at large spatial scales^23^. Thus, variations in microbial biogeography, in particular, deviations from microbial biomes as defined by the CSH, can potentially be used in monitoring environmental changes, and might therefore be valuable tools in marine management.

## Methods

Occupancy-frequency distributions were analyzed as described in^20^. In brief, an equivalent to Tokeshi’s test of bimodality was performed using Mitchell-Olds’ and Shaw’s test^22^ for the location of quadratic extremes. The global ocean transect data were downloaded from the European Nucleotide Archive (ENA; accession number PRJEB7988) and the NCBI Sequence Read Archive (SRA; accession number SRP031469), Tara Oceans and Malaspina transects, respectively. For *in silico* tests we first tested the effect of sequencing depth and subsampled the Tara oceans dataset to 100,000 sequence reads and rarefied this subsampled dataset to 1000, 2000, 5000, 10,000, 20,000, 40,000, 60,000 and 80,000 sequence reads and plotted the cumulative number of satellite compared to core populations. We further created one randomized dataset by shuffling the presence/absence of OTUs using the “permatfull” function in R by keeping the sample sums (100,000 sequence reads) static and performing 999 unrestricted permutations of the OTUs from the subsampled Tara oceans dataset and picked one random permuted community as mock community and rarefied as above. Thirdly, for the occupancy-frequency analyses we subsampled the Tara oceans and Malaspina dataset to 40,590 and 25,000 sequence reads, respectively, well above the suggested lower limit for diversity analyses of marine microbial assemblages^23^ and the 25,000 sequence reads noted in the test above (Fig. S2). Finally, we randomized each of the subsampled Tara oceans and Malaspina dataset and specific oceanic regions (North Atlantic, South Pacific and South Atlantic, Brazil basin, Tara oceans and Malaspina datasets respectively) and we used one randomized dataset as a mock community and this artificial community were randomized as described above. All statistical tests were performed in R 3.3.3^24^, and using the package “Vegan”^25^.

## Supporting information

### Supplementary figures and tables

**Figure S1.**
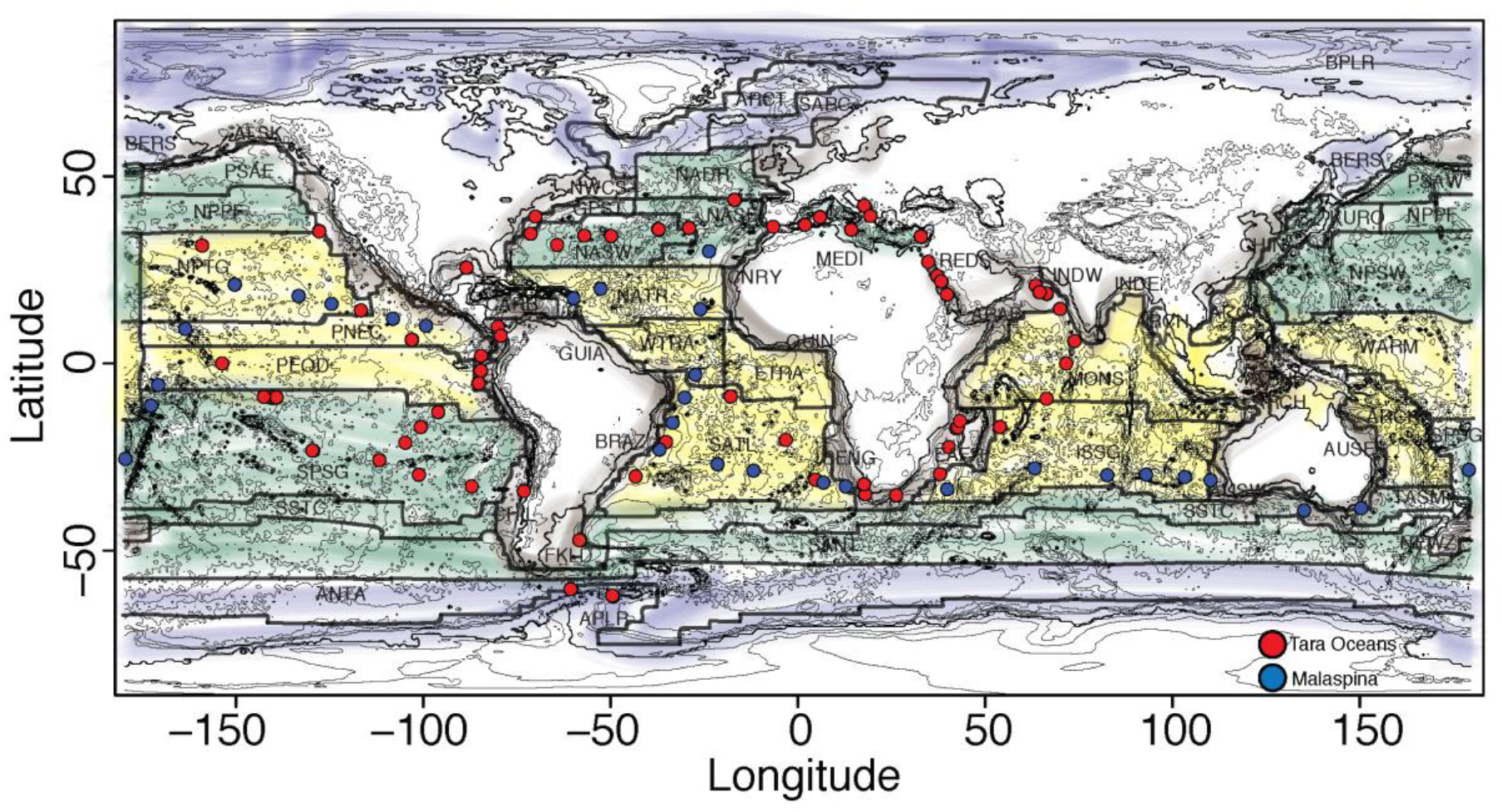
Schematic map of sampled stations in the Tara oceans^6^ and Malaspina^7^ datasets included in the analyses and superimposed with Longhurst’s biogeographical division of the ocean into biomes and provinces. Blue, Brown, Green and Yellow color denote Polar, Coastal, Westerlies and Trades biomes, respectively.

**Figure S2.**
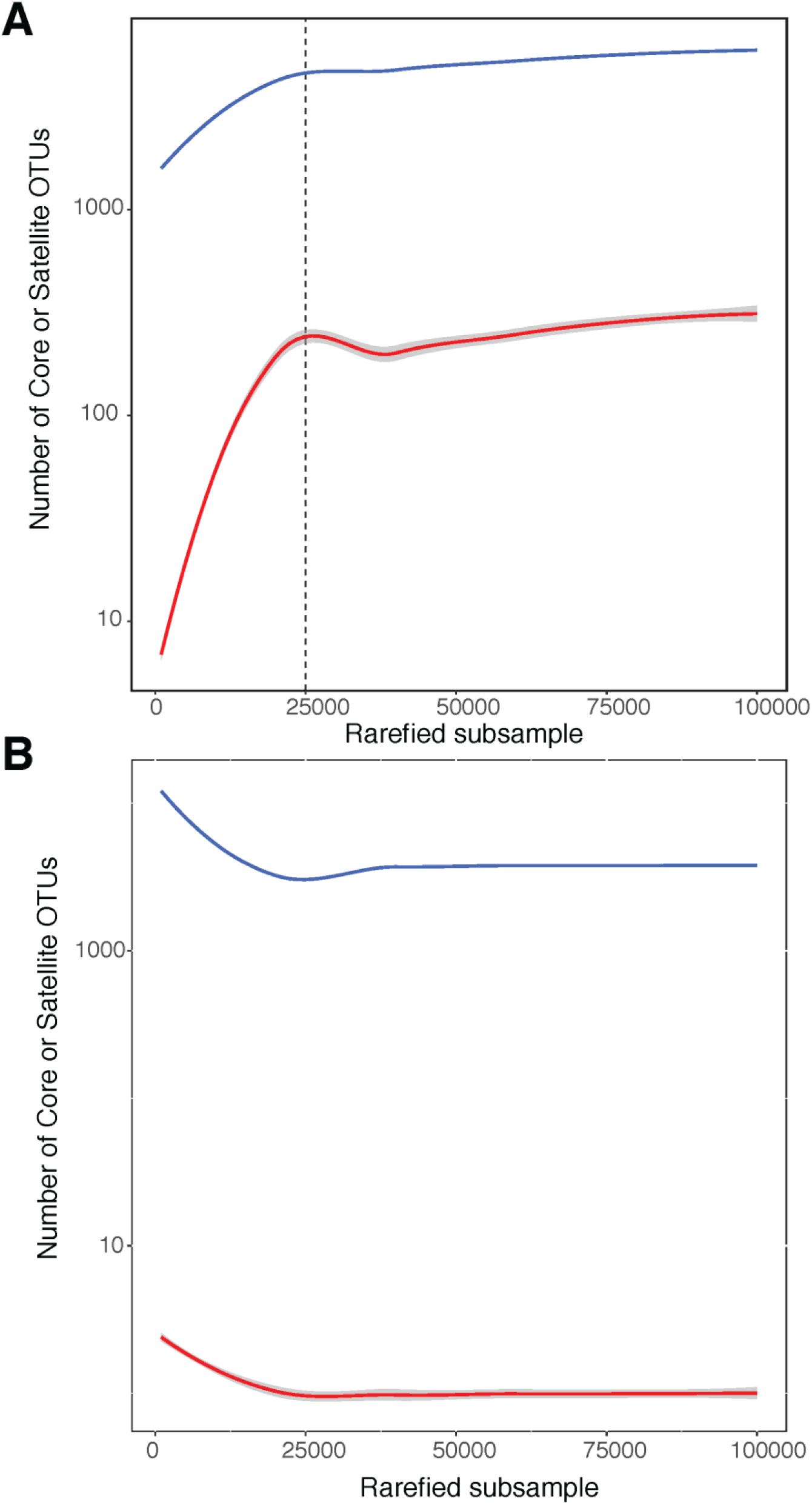
*In silico* tests of the effects of sequencing depth on the cumulative number of core compared to satellite populations for the rarefied total community obtained from Tara Oceans (A), and one mock community (B) obtained from randomizing the Tara Oceans dataset and rarefying as in (A).

